## Supplementary Table and Figures for "Structure of the SARS-CoV NSP12 polymerase bound to NSP7 and NSP8 co-factors"

| | NSP7-NSP8-NSP12 | NSP8-NSP12 |
|---|---|---|
| EMDB | EMD-0520 | EMD-0521 |
| **Data and reconstruction statistics** | | |
| Microscope | Talos Actica | Talos Actica |
| Voltage (kV) | 200 | 200 |
| Detector | Gatan K2 Summit | Gatan K2 Summit |
| Dose rate ($e^-$/pix/sec) | 5.69 | 5.69 |
| Exposure (s) | 11.75 | 11.75 |
| Dose ($e^-$/Å$^2$) | 50.5 | 50.5 |
| Frames | 47 | 47 |
| Defocus Range (μm) | 0.4 – 1.0 | 0.4 – 1.0 |
| Initial Particles | 609,107 | 609,107 |
| Final Particles | 71,046 | 71,262 |
| B-factor (Å$^2$) | -49 | -64 |
| Resolution (Å) | 3.1 | 3.5 |
| **Coordinate model refinement** | | |
| PDB | 6NUR | 6NUS |
| Residues | 1,087 | 827 |
| RMSD Bonds (Å) | 0.016 | 0.016 |
| RMSD Angles (˚) | 1.56 | 1.58 |
| Ramachandran | | |
| Favored (%) | 98.6 | 98.8 |
| Allowed (%) | 1.4 | 1.2 |
| Outliers (%) | 0.0 | 0.0 |
| Rotamer Outliers (%) | 0.5 | 0.1 |
| Clash score | 2.04 | 0.92 |
| Molprobity score | 0.97 | 0.78 |
| EM Ringer Score | 4.15 | 2.07 |

**Supplementary Table 1. Data collection and refinement.** Resolution was estimated using a gold-standard 0.143 FSC cutoff. Coordinate model quality was assessed using Molprobity[1] and EMRinger[2].

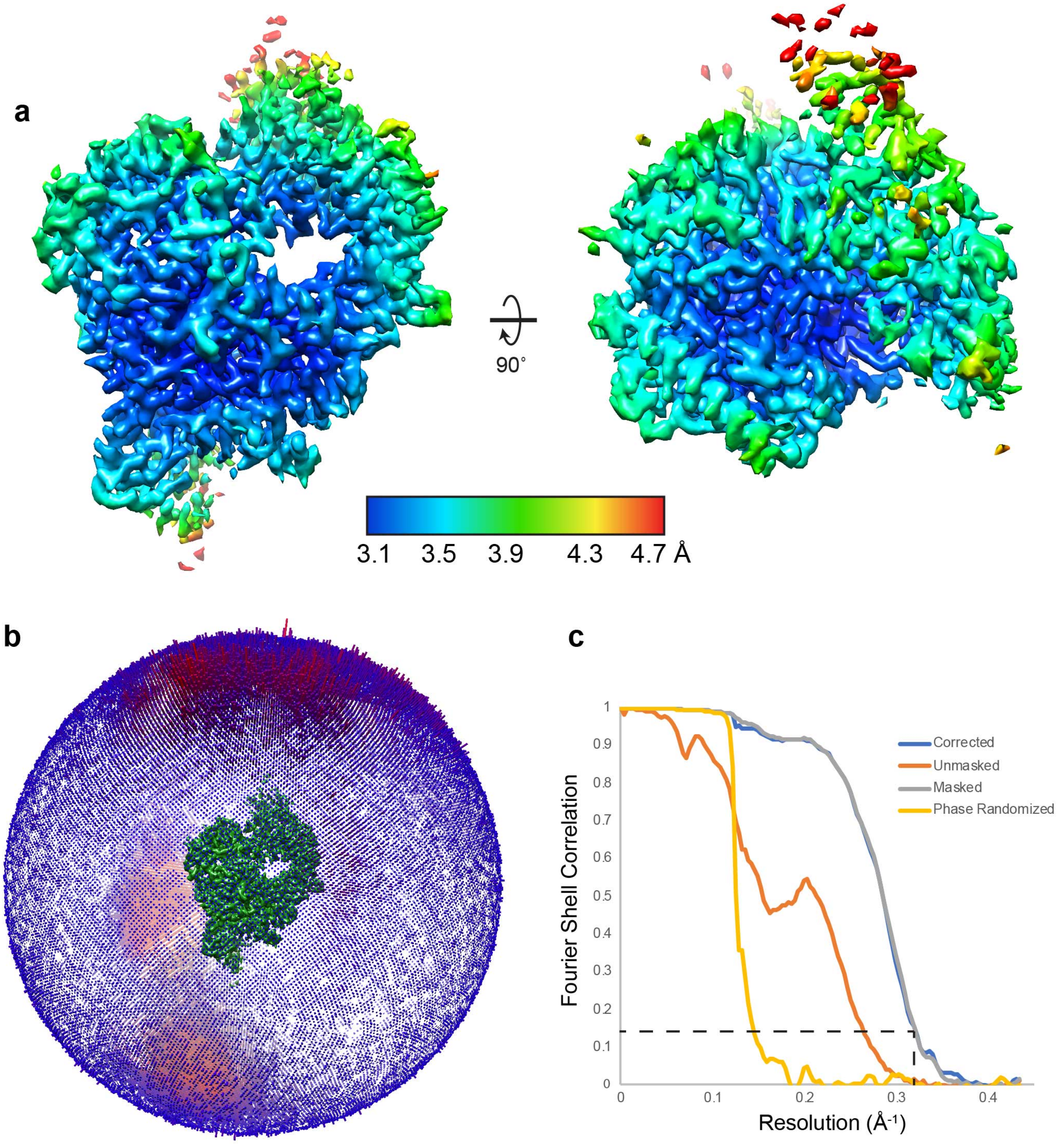

**Supplemental Fig 1. NSP7-NSP8-NSP12 cryoEM map validation.** Local resolution estimation, angular distribution and FSC curves were calculated in RELION-3.0[3].

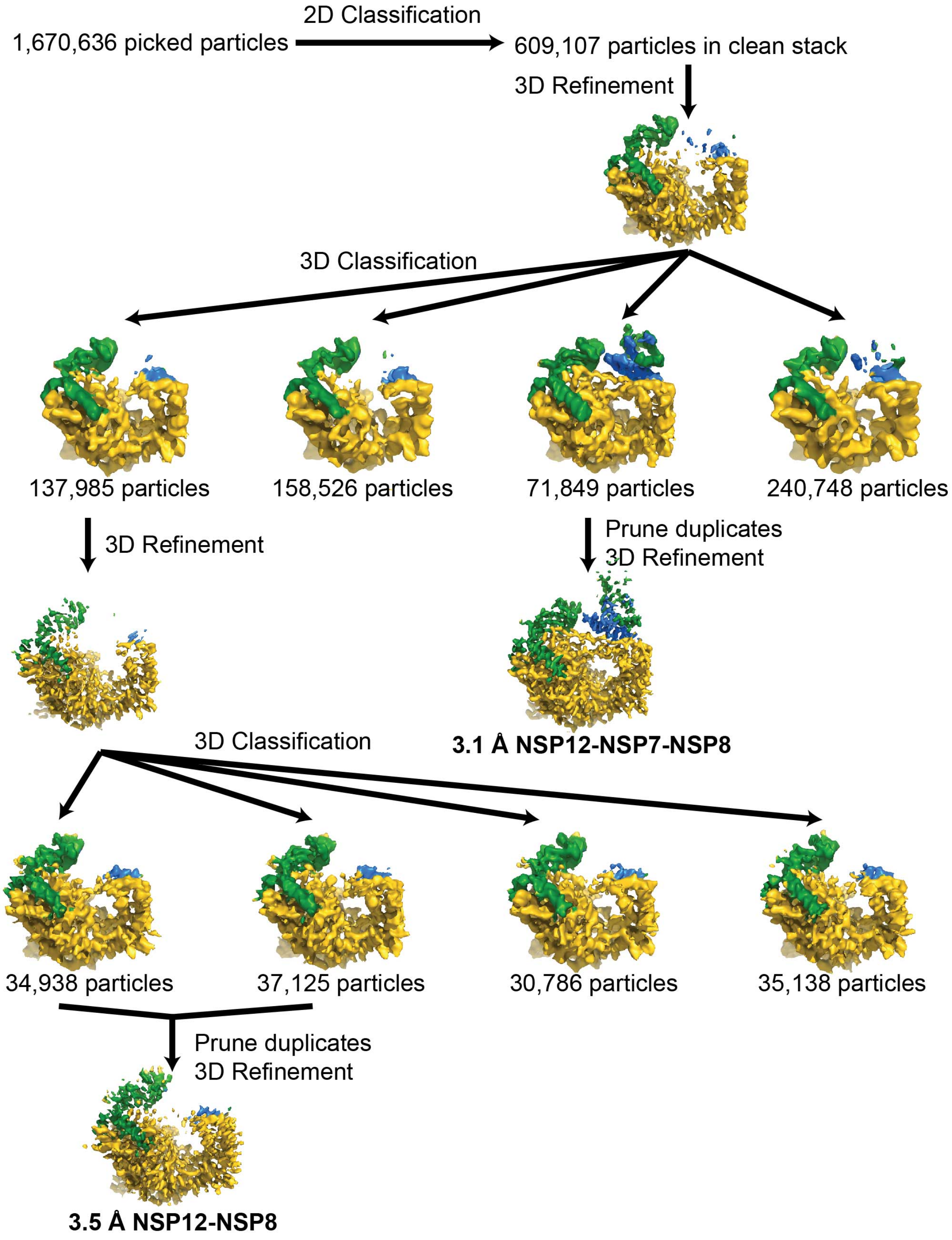

**Supplementary Figure 2: Classification and refinement of cryoEM data.**

```
PDCV    -----SLQNSAYLNRVTG-SSDARLEPLQPGTQPDAVKRAFHVHN--DTTSGIFLSTKSN   52
NL63    ----------SYLNRARG-SSAARLEPCN-GTDIDKCVRAFDIYN--KNVSFLGKCLKMN   46
IBV     SAAGAPDFDKNYLNRVRG-SSEARLIPLANGCDPDVVKRAFDVCN--KESAGMFRNLKRN   57    NiRAN
MHV     ------SKDTNFLNRVRGTSVNARLVPCASGLDTDVQLRAFDICN--ANRAGIGLYYKVN   52
SARS    -----SADASTFLNRVCG-VSAARLTPCGTGTSTDVVYRAFDIY--NEKVAGFAKFLKTN   52
MERS    ------SKDSNFLNRVRGSIVNARIEPCSSGLSTDVVFRAFDICNYKAKVAGIGKYYKTN   54
                 :***. *     **: *    *  . *    ***.:       : :    * *

PDCV    CARFKTTRSALPLPNKGEVELYFVTKQCAAKVFEIEEECYNAISTELYTTDDTFGVLAKT   112
NL63    CVRFKNAD---------LKDGYFVIKRCTKSVMEHEQSMYNLLNF--------SGALAEH   89
IBV     CARFQEVRDTE-DGNLEYCDSFFVVKQTTPSNYEHEKSCYEDLKS---------EVTADH   107   NiRAN
MHV     CFRFQRVDEEG-----NKLDKFFVVKRTNLEVYNKEKECYELTKD--------CGVVAEH   99    Motif A_N
SARS    CCRFQEKDEEG-----NLLDSYFVVKRHTMSNYQHEETIYNLVKD--------CPAVAVH   99
MERS    TCRFVELDDQG-----HHLDSYFVVKRHTMENYELEKHCYDLLRD--------CDAVAPH   101
          **                : :** *:    .   : *:   *:              . *

                              ↓First visible amino acid
PDCV    EFFKFD----KIPNVNRQYLTKYTLLDLAYALRHLSTS-KDVIQEILITMCGTP---EDW   164
NL63    DFFTWKDGRVIYGNVSRHNLTKYTMMDLVYAMRNFDEQNCDVLKEVLVLTGCCDNS----   145
IBV     DFFVFNK---NIYNISRQRLTKYTMMDFCYALRHFDPKDCEVLKEIFVTYGCIEDYHPKW   164   NiRAN
MHV     EFFTFDVEGSRVPHIVRKDLSKFTMLDLCYALRHFDRNDCSTLKEILLTYAECDES----   155   Motif B_N
SARS    DFFKFRVDGDMVPHISRQRLTKYTMADLVYALRHFDEGNCDTLKEILVTYNCCDDD----   155
MERS    DFFIFDVDKVKTPHIVRQRLTEYTMMDLVYALRHFDQN-SEVLKAILVKYGCCDVT----   156
        :** :          :: *: *:::*: *: **:*::.     ..:: :::

PDCV    F--GENWFDPIENPSFYKEFHKLGDILNRCVLNANKFASACIDAGLVGILTPDNQDLLGQ   222
NL63    YFDSKGWYDPVENEDIHRVYASLGKIVARAMLKCVALCDAMVAKGVVGVLTLDNQDLNGN   205
IBV     FEENKDWYDPIENPKYYAMLAKMGPIVRRALLNAIEFGNLMVEKGYVGVVTLDNQDLNGK   224   NiRAN
MHV     YFQKKDWYDFVENPDIINVYKKLGPIFNRALLNTANFADTLVEAGLVGVLTLDNQDLYGQ   215   Motif C_N
SARS    YFNKKDWYDFVENPDILRVYANLGERVRQSLLKTVQFCDAMRDAGIVGVLTLDNQDLNGN   215
MERS    YFENKLWFDFVENPSVIGVYHKLGERVRQAILNTVKFCDHMVKAGLVGVLTLDNQDLNGK   216
        :   *:* :**  .       .:*   . :.:*:    :  .      *  **::* ***** *:

PDCV    IYDFGDFIITQPGNGCVDLASYYSYLMPIMSMTHMLKCECMDS----DGNPLEYDGFQYD   278
NL63    FYDFGDFVVSLPNMGVPCCTSYYSYMMPIMGLTNCLASECFVKSDIFGSDFKTFDLLKYD   265
IBV     FYDFGDFQKTALGAGVPVFDTYYSYMMPIIAMTDALAPERYFEYDVH-KGYKSYDLLKYD   283   NiRAN, Interface
MHV     WYDFGDFVKTVPCCGVAVADSYYSYMMPMLTMCHALDSELFV----N-GTYREFDLVQYD   270   Motif C_N
SARS    WYDFGDFVQVAPGCGVPIVDSYYSLLMPILTLTRALAAESHMDADLA-KPLIKWDLLKYD   274
MERS    WYDFGDFVITQPGSGVAIVDSYYSYLMPVLSMTDCLAAETHRDCDFN-KPLIEWPLTEYD   275
         ******       *      :*** :**:: :        *  *              :    :**

                                  ●    ●   ●   ●                      Zinc Coordinating
PDCV    FTDFKLGLFEKYFKYWDRPYHPNTVECPDDRCVLHCANFNVLFAMCIPNTAFGNLCSRAT   338
NL63    FTEHKENLFNKYFKHWSFDYHPNCSDCYDDMCVIHCANFNTLFATTIPGTAFGPLCRKVF   325
IBV     YTEEKQEMFQKYFKYWDQEYHPNCRDCSDDRCLIHCANFNILFSTLIPQTSFGNLCRKVF   343   Interface
MHV     FTDFKLELFNKYFKHWSMTYHPNTSECEDDRCIIHCANFNILFSMVLPKTCFGPLVRQIF   330
SARS    FTEERLCLFDRYFKYWDQTYHPNCINCLDDRCILHCANFNVLFSTVFPPTSFGPLVRKIF   334
MERS    FTDYKVQLFEKYFKYWDQTYHANCVNCTDDRCVLHCANFNVLFAMTMPKTCFGPIVRKIF   335
        :*: :  :*::***:*:.  ** *  :* ** *::****** **:  :* *.** :   :

PDCV    VDGHLVVQTVGVHLKELGIVLNQDVTTHMANINLNTLLRLVGDPTTIASVSDKCVDLRTP   398
NL63    IDGVPLVTTAGYHFKQLGLVWNKDVNTHSVRLTITELLQFVTDPSLIIASSPALVDQRTI   385
IBV     VDGVPFIATCGYHSKELGVIMNQDNTMSFSKMGLSQLMQFVGDPALLVGTSNNLVDLRTS   403   Interface
MHV     VDGVPFVVSIGYHYKELGVVMNMDVDTHRYRLSLKDLLLYAADPALHVASASALLDLRTC   390
SARS    VDGVPFVVSTGYHFRELGVVHNQDVNLHSSRLSFKELLVYAADPAMHAASGNLLLDKRTT   394
MERS    VDGVPFVVSCGYHYKELGLVMNMDVSLHRHRLSLKELMMYAADPAMHIASSNAFLDLRTS   395
        :**   .: : * * ::**::* *      .: :. *:  . **:   .  .    :* **

PDCV    CQTLATMSSGIAKQSVKPGHFNQHFYKHLLDSNLLD-QLGIDMRHFYYMQDGEAAITDYS   457
NL63    CFSVAALSTGLTNQVVKPGHFNEEFYNFLRLRGFFDEGSELTLKHFFFAQNGDAAVKDFD   445
IBV     CFSVCALASGITHQTVKPGHFNKDFYDFAEKAGMFKEGSSIPLKHFFYPQTGNAAINDYD   463   Interface, Fingers
MHV     CFSVAAITSGVKFQTVKPGNFNQDFYEFILSKGLLKEGSSVDLKHFFFTQDGNAAITDYN   450
SARS    CFSVAALTNNVAFQTVKPGNFNKDFYDFAVSKGFFKEGSSVELKHFFFAQDGNAAISDYD   454
MERS    CFSVAALTTGLTFQTVRPGNFNQDFYDFVVSKGFFKEGSSVTLKHFFFAQDGNAAITDYN   455
        * :*:.:::..:  * *:**:**:.**..      .::.    : ::**:: * *:**:.*:.

                                          ●                          Zinc Coordinating
PDCV    YYRYNTPTMVDIKMFLFCLEVADKYLEPYEGGCINAQSVVVSNLDKSAGYPFNKLGKARN   517
NL63    FYRYNKPTILDICQARVTYKIVSRYFDIYEGGCIKACEVVVTNLNKSAGWPLNKFGKASL   505
IBV     YYRYNRPTMFDIRQLLFCLEVTSKYFECYEGGCIPASQVVVNNLDKSAGYPFNKFGKARL   523   Fingers
MHV     YYKYNLPTMVDIKQLLFVVEVVNKYFEIYEGGCIPATQVIVNNYDKSAGYPFNKFGKARL   510   Motif G
SARS    YYRYNLPTMCDIRQLLFVVEVVDKYFDCYDGGCINANQVIVNNLDKSAGFPFNKWGKARL   514
MERS    YYSYNLPTMCDIKQMLFCMEVVNKYFEIYDGGCLNASEVVVNNLDKSAGHPFNKFGKARV   515
        :* ** **: **        .   ::..:*:: *:***: * .*:*.* :**** *:** ***
```

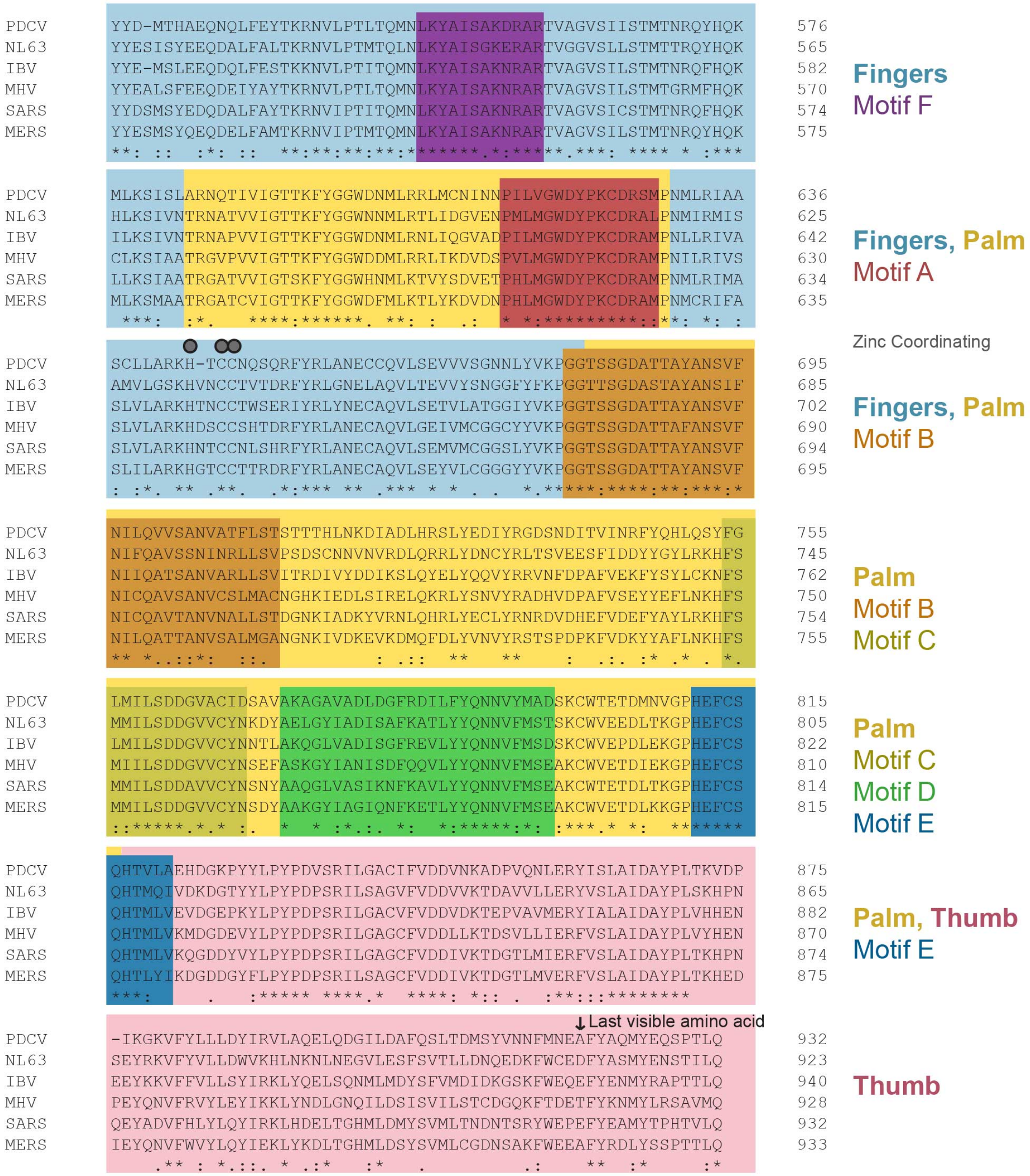

**Suplemental Figure 3: Annotated sequence alignment of coronavirus NSP12.** Sequences of porcine deltacoronavirus (PDCV), human coronavirus NL63, infectious bronchitis virus (IBV), murine hepatitis virus (MHV), human SARS coronavirus and MERS coronavirus NSP12 were aligned with Clustal Omega[4] and annotated by functional region[5-8] and structural observations noted in the main text.

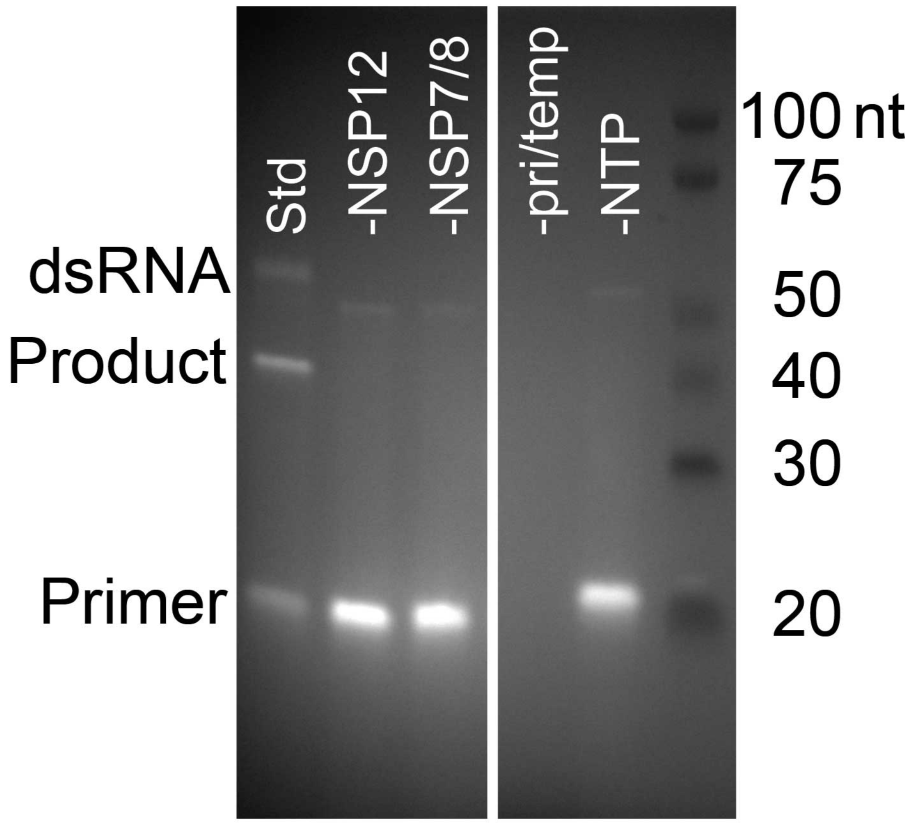

**Supplementary Figure 4: Primer extension activity of the SARS-CoV NSP7-NSP8-NSP12 complex.** A 20 nt fluorescently labeled primer was extended on a 40 nt template and analyzed by denaturing TBE-Urea PAGE. Negative controls lacking NSP12, NSP7 and NSP8, the annealed primer/template and NTPs are indicated. Molecular size markers are indicated on the right. Some extended and unextended double stranded RNAs which are the result of incomplete denaturation are also visible in the range of 50-60 nt.

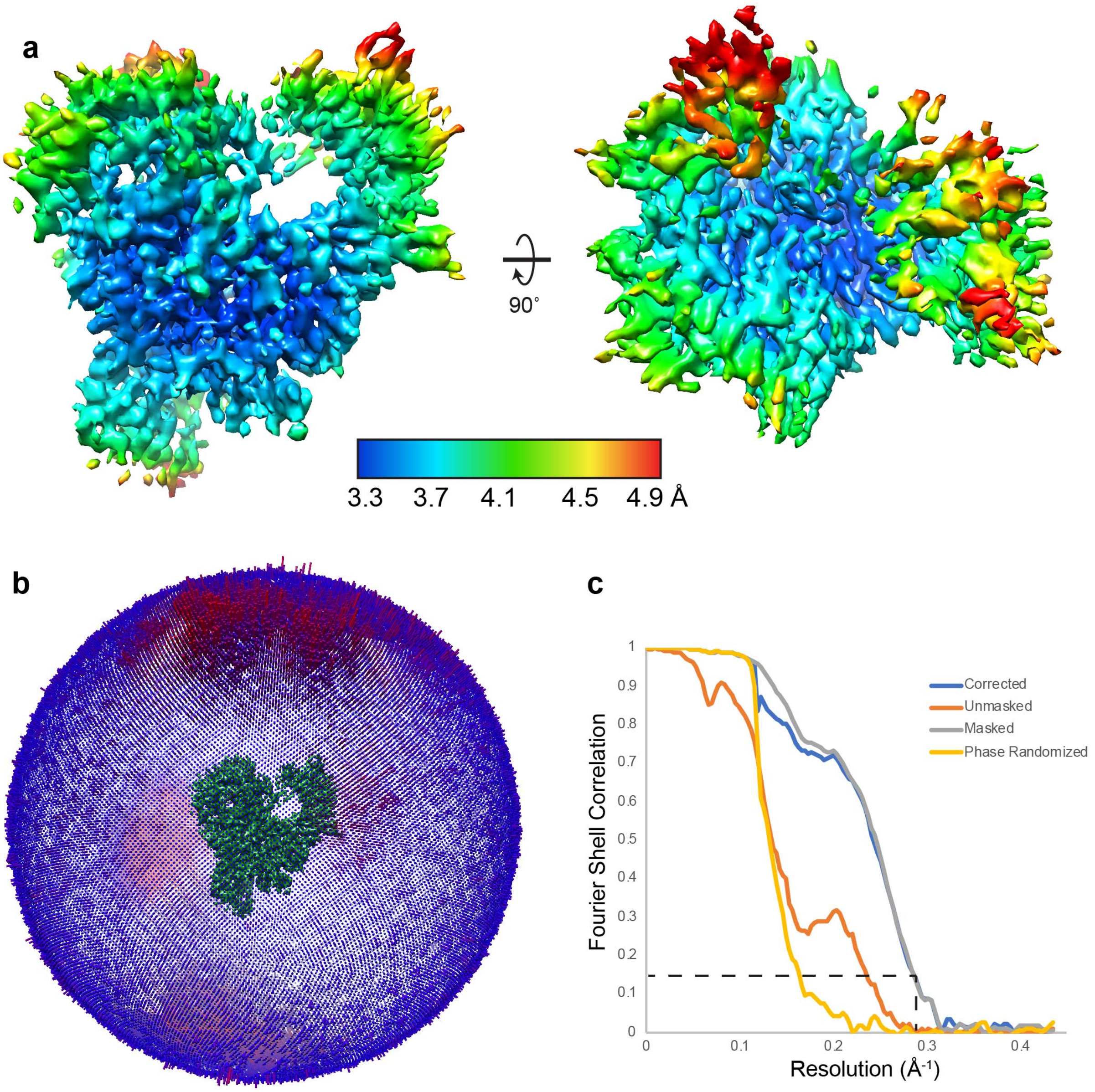

**Supplemental Fig 5. NSP8-NSP12 cryoEM map validation.** Local resolution estimation, angular distribution and FSC curves were calculated in RELION-3.0[3].

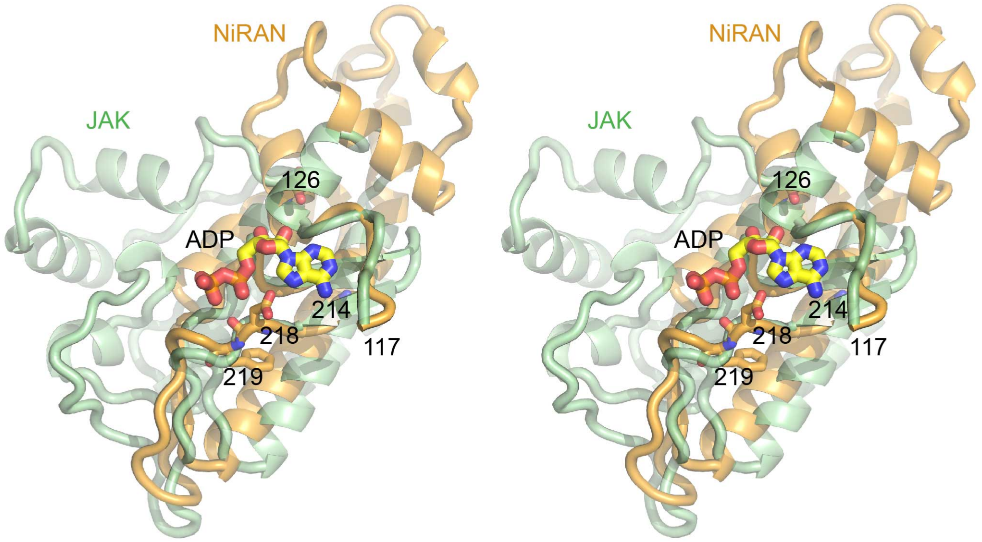

**Supplementary Fig. 6: The NSP12 NiRAN domain contains structural homology to the C-terminal large domain of kinases.** Stereoview of a superposition of the SARS-CoV NiRAN domain with the C-terminal kinase domain of human JAK (6C7Y.pdb) demonstrates similar structures neighboring the JAK nucleotide binding site. Nidovirus conserved residues Asp126, Gly214, Asp218 and Phe219 are labeled and shown as sticks. A GDP nucleotide bound in the JAK kinase domain structure is also shown.

### Supplementary References

1 Williams, C. J. *et al.* MolProbity: More and better reference data for improved all-atom structure validation. *Protein Sci* **27**, 293-315, (2018).

2 Barad, B. A. *et al.* EMRinger: side chain-directed model and map validation for 3D cryo-electron microscopy. *Nat Methods* **12**, 943-946, (2015).

3 Zivanov, J. *et al.* New tools for automated high-resolution cryo-EM structure determination in RELION-3. *Elife* **7**, (2018).

4 Sievers, F. & Higgins, D. G. Clustal Omega for making accurate alignments of many protein sequences. *Protein Sci* **27**, 135-145, (2018).

5 Bruenn, J. A. A structural and primary sequence comparison of the viral RNA-dependent RNA polymerases. *Nucleic Acids Res* **31**, 1821-1829, (2003).

6 Gorbalenya, A. E. *et al.* The palm subdomain-based active site is internally permuted in viral RNA-dependent RNA polymerases of an ancient lineage. *J Mol Biol* **324**, 47-62, (2002).

7 Lehmann, K. C. *et al.* Discovery of an essential nucleotidylating activity associated with a newly delineated conserved domain in the RNA polymerase-containing protein of all nidoviruses. *Nucleic Acids Res* **43**, 8416-8434, (2015).

8 Poch, O., Sauvaget, I., Delarue, M. & Tordo, N. Identification of four conserved motifs among the RNA-dependent polymerase encoding elements. *Embo j* **8**, 3867-3874, (1989).
